# EEG Alpha Power and Pupil Diameter Reflect Endogenous Auditory Attention Switching and Listening Effort

**DOI:** 10.1101/2021.07.29.453646

**Authors:** Stephanie Haro, Hrishikesh M. Rao, Thomas F. Quatieri, Christopher J. Smalt

## Abstract

Auditory attention describes a listeners focus on an acoustic source while they ignore other competing sources that might be present. In an environment with multiple talkers and background noise (i.e. the cocktail party effect), auditory attention can be difficult, requiring the listener to expend measurable cognitive effort. A listener will naturally interrupt sustained attention on a source when switching towards another source during conversation. This change in attention is potentially even more taxing than maintaining sustained attention due to the limits of human working memory, and this additional effort required has not been well studied. In this work, we evaluated an attention decoder algorithm for detecting the change in attention and investigated cognitive effort expended during attentional switching and sustained attention. Two variants of endogenous attention switching were explored: the switches either had in-the-moment decision making or a pre-defined attentional switch time. A least-squares, EEG-based, attention decoding algorithm achieved 64.1% accuracy with a 5-second correlation window and illustrated smooth transitions in the attended talker prediction through switches in sustained attention at approximately half of the analysis window size (2.2 seconds). The expended listening effort, as measured by simultaneous electroencephalography (EEG) and pupillometry, was also a strong indicator of switching. Specifically, centrotemporal alpha power [F(2, 18) = 7.473, P = 0.00434] and mean pupil diameter [F(2, 18) = 9.159, P = 0.0018] were significantly different for trials that contained a switch in comparison to sustained trials. We also found that relative attended and ignored talker locations modulate the EEG alpha topographic response. This alpha lateralization was found to be impacted by the interaction between experimental condition and whether the measure was computed before or after the switch [F(2,18) = 3.227, P = 0.0634]. These results suggest that expended listening effort is a promising feature that should be pursued in a decoding context, in addition to speech and location-based features.

## 1. Introduction

Real-world listening situations often contain multiple competing talkers and listeners must engage auditory attention to focus onto one source. In voluntary sustained attention, an *endogenous* process, the listener applies salience towards a source and top-down mechanisms influence how the source is represented in the cortex (Posner et al., 1984; Golumbic et al., 2013). In most environments, however, listeners do not sustain their attention to one talker continuously. Listeners may switch their attention endogenously, shifting their attention between sources at their discretion. Meanwhile, sources also vie to *exogenously* capture listener attention, employing bottom-up processes once successful (Posner et al., 1984).

Exogenous and endogenous attention are distinct yet intertwined in audition. Distinct brain regions have been shown to be involved in endogenous auditory attention (Hill & Miller, 2010; Lee et al., 2013; Larson & Lee, 2014). Their studies have focused on characterizing attention between location and pitch, two core features that differ between sources in cocktail-party scenarios. The frontal-parietal region was found to be activated during endogenous auditory attention towards sources that differ in both space and pitch (Hill & Miller, 2010). Next, the frontal eye field region was found to be activated in preparation for and during endogenous attention towards sources (Lee et al., 2013). A more temporally resolved modality was used to find distinct parietal activations during endogenous switches (Larson & Lee, 2014). The right temporoparietal junction (RTPJ) and the left inferior parietal supramarginal part (LIPSP) were active during switches between sources that differed in space and pitch respectively. Exogenous distractor stimuli have also been found to counter endogenous attention’s enhancement effects in an auditory study of natural soundscapes (Huang & Elhilali, 2020). The previously mentioned attention results have been determined using small speech tokens such as alphabetic characters, unrelated sentences from a corpus, or non-speech scene stimuli (Hill & Miller, 2010; Lee et al., 2013; Larson & Lee, 2014; Huang & Elhilali, 2020). The regions involved with switching attention naturally between continuous speech sources have yet to be characterized and will likely involve brain regions beyond those previously mentioned.

Real-world attention occurs in scenes that are more complex than the stimuli used in conventional clinical hearing assessments. These real scenes often involve multiple speech sources and reverberation that recruit speech-specific auditory processes (Liberman et al., 2016). This complexity may provide a suite of cues that can be leveraged by the listener during auditory attention. A longer listening task may lead to more opportunities to latch attention and greater overall comprehension due to the continuous speech context. Identifying regions engaged in switching can potentially be used to track attention states. These states can then be used to control stimuli enhancement which can improve the listener’s experience. Altering the relative levels of attended and ignored stimuli can reduce listening effort and enhance attended stimuli entrainment (Seifi Ala et al., 2020; Presacco et al., 2019; Mirkovic et al., 2019). Speech enhancement has the capacity to improve listener quality of life in individuals of all ages and levels of hearing loss (Griffin et al., 2019; Liberman et al., 2016; Ciorba et al., 2012; Griffin et al., 2019).

Auditory attention decoding (AAD) describes the process of using cortical recordings to identify to whom a listener is attending when multiple talker sources are competing for the listeners attention. AAD in combination with speaker separation has the potential to be incorporated into cognitively-controlled hearing aids to provide auditory enhancement in speech-rich scenes that traditional hearing aids struggle with (Popelka & Moore, 2016; Borgström et al., 2021). The majority of these studies’ protocols ask listeners to sustain attention, not invoking endogenous and exogenous switches in attention (Geirnaert et al., 2021). However, it is critical to study both types of attention switching given the prevalence of switching in real-world conditions. Various speech features and cortical recording modalities have been used to encode and decode attended stimuli (Mesgarani & Chang, 2012; O’Sullivan et al., 2015; OSullivan et al., 2017; Ciccarelli et al., 2019; Ding & Simon, 2012; Akram et al., 2016; Puvvada & Simon, 2017). These decoding algorithms have relied on reactive decoding of the already attended stimuli, creating a lag in the enhancement. Endogenous switches are associated with top-down attentional preparatory activity in contrast to exogenous attention switches (Lee et al., 2013), and thus tracking endogenous preparatory activity might aid in faster enhancement in comparison to reactive decoding. Identifying preparatory features that accompany attention switches could aid in more robust auditory enhancement when combined with attended stimuli decoding.

Recent work has begun incorporating exogenous and endogenous switches in their attention decoding protocols to explore alternative attention modeling techniques (Akram et al., 2016; Teoh & Lalor, 2019; Miran et al., 2018, 2020). In some auditory switching studies, the switch time is determined by the protocol (Hill & Miller, 2010; Lee et al., 2013; Larson & Lee, 2014; Akram et al., 2016; Teoh & Lalor, 2019). These studies direct attention switching using acoustic cues - directing the listener to switch sources when a gap in the stimulus occurs (Akram et al., 2016) or instructing the listener to switch attended talker location in order to track a dynamic talker (Teoh & Lalor, 2019). However when naturalistic endogenous attention switches are studied, the true switch time is known by the listener and must be extracted. For example, a button press has been used to record endogenous switch time (Miran et al., 2018, 2020). Unfortunately, a button press may create switch-locked pre-motor planning and muscle artifacts in the data, potentially confounding endogenous switching feature interpretation (Johari et al., 2019; Stephen, 2019).

In this study, we investigated endogenous switches of sustained attention between competing multi-talker sources. Listeners were asked to remember when they endogenously switched using a clock in order to remove an explicit evoked response, e.g. a motion artifact from a button press. For the first analysis, we performed regularized least-squares decoding of the attended talker envelope (Crosse et al., 2016). We demonstrate decoder prediction behavior on data that contains attention switches and additional higher-order processes of memorization and decision making that are incorporated into the protocol. Next, we quantified the effort involved with endogenous switching using measures of EEG alpha power and pupil diameter which proved successful in characterizing listening effort during sustained attention (Seifi Ala et al., 2020). Lastly, we analyzed alpha power activity related to the relative locations of the attended and unattended talker locations which has been characterized during attention between spatially separated syllable streams (Deng et al., 2020).

## 2. Methods

### 2.1. Experimental Protocol

Ten native English speakers (5F, 5M), with self-reported normal hearing, participated in this study. They provided informed consent to an experimental protocol that was approved by the MIT Committee on the Use of Humans as Experimental participants and the The U.S. Army Medical Research and Development Command (USAMRDC), Human Research Protection Office (HRPO). Participants were asked to sit in a sound treated booth between two loudspeakers positioned 6 feet away at +/-45 degrees. The left and right loudspeakers presented male talkers reading Twenty Thousand Leagues Under the Sea and Journey to the Center of the Earth audiobooks, respectively (O’Sullivan et al., 2019). We simultaneously recorded participant EEG and pupillometry using a dry electrode EEG system (Wearable Sensing DSI-24) and eye tracking glasses (SMI ETG2), respectively. The EEG and pupillometry data were sampled at 300Hz and 120 Hz, respectively. A monitor situated in front of the participant displayed various stages of the protocol.

Figure 1 diagrams the instructional stages of a trial on the left of the figure and the time course of the three experimental conditions on the right. This protocol consisted of 60 one-minute trials (20 trials of each experimental condition). At the beginning of a trial, the screen presented the trial task. Each trial task was defined as a combination of the experimental condition and initial attended talker (left or right). Each experimental condition had an equal number of trials that began with either left or right talker attention. The experimental condition presentation order was randomized and determined using MATLAB’s uniformly distributed pseudorandom integer generator. During a trial, after approximately 30 seconds of attention towards the initial attended talker, listeners performed one of three tasks. Depending on the indicated experimental condition, listeners were to switch attention at their own discretion (at-will), switch attention at a directed time (directed), or not switch attention at all (sustained).

**Figure 1:**
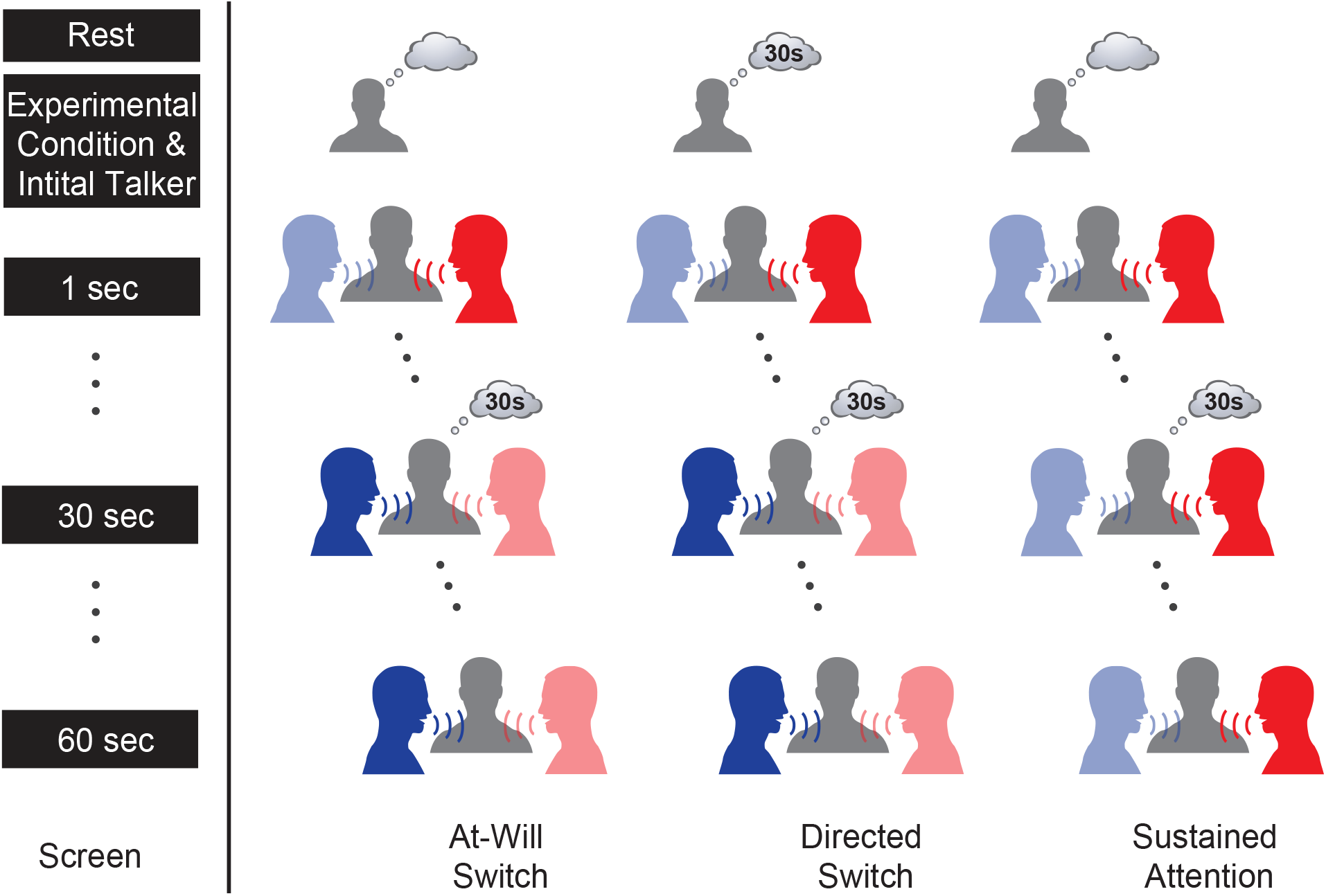
Auditory attention protocol experimental conditions compared. During the protocol, listeners were presented with two competing spatially separated audiobook stimuli. They were asked to begin each trial attending to one talker and ignoring the other. For two of the experimental conditions, listeners were asked to switch talkers approximately halfway through the trial. Listeners either switched attention at their discretion (at-will switch) or switched their attention at a time specified before the trial began (directed switch). In the third experimental condition, listeners were asked to keep their attention on the initial talker for the whole trial (sustained attention). At the end of the trial, listeners recalled the time event they either switched or continued to sustain attention.

To record the attention switch without the use of button press, we incorporated a time memorization task into all three experimental conditions. From the onset of the trial, the screen presented the elapsed time, updated once a second. The at-will switch involves an on-demand, listener-initiated switch; the listener used the clock to mentally note when they switched at their discretion. The directed switch involved the listener switching at a time specified before the trial began. The participant used the clock to perform the attention switch at the pre-determined time. Note that the directed switch lacks the added online decision-making task of when to switch. The directed condition may not resemble a realistic endogenous switch but serves as a reference, for comparison with the at-will switch. For the sustained condition, the listener attended to the same talker for the whole trial but was tasked to remember a time only once they saw it on the clock. This time task is modeled after online decision-making that would occur in an endogenous switch of attention, but without the actual switch. The sustained experimental condition controlled for the executive functioning tasks used in the at-will experimental condition (decision making and remembering time).

Figure 1 illustrates the listener memory state across the experimental conditions in the thought bubbles. All three conditions’ timing events were only permitted to occur between [25,35] seconds in order to ensure ample data before and after the switch. The directed switch time was randomly generated. For the other experimental conditions, participants were instructed to randomize their timing events between the range of [25,35] seconds themselves. For the rest of the analysis, all trial times were normalized relative to the timing event as time 0 seconds. The clock updated once per second instead of a finer resolution in order to prevent increased visual processing load and reduce the complexity of the time memorization task. Any physiological measures seen around the normalized time of 0 seconds can be attributed to executive functioning related to the listener’s decision making, committing the switch time to memory, and/or switching auditory attention between sources. Between each trial, participants recalled the trial’s timing event and answered two 4-choice comprehension questions using a wireless gaming controller. Each of the 10 participant collections contain 60 minutes worth of EEG and pupillometry data as well as two comprehension responses for each trial. The protocol contains characteristics that should elicit measurable effort such as a reasonably difficult task that provides listener engagement and motivation (Winn et al., 2018). Listeners attend to continuous audio book speech stimuli which keeps their engagement throughout the experiment more than other simpler stimuli would. The act of attending to a source while ignoring another is reasonably difficult but not too taxing. Participants were motivated to value attending to the proper talker at all time since they need to answer comprehension questions from both halves of the trial. We believe participants were motivated to heed the three experimental conditions tasks equally.

### 2.2. Auditory Attention Decoding

We used a regularized least-squares decoding approach to predict to whom the listener was attending to before, during, and following shifts in auditory attention induced by our protocol. Least-squares decoding was used to transform a window of EEG signal into an attended talker envelope speech prediction, 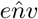, using a linear weight matrix mapping (Crosse et al., 2016). This method operates on the basis of the attended talker speech envelope being encoded more robustly in the listener’s EEG than the ignored talker’s speech envelope (Golumbic et al., 2013; O’Sullivan et al., 2015). Pre-processing was kept at a minimum because decoding that limits pre-processing is faster and therefore a better fit for real-time processing applications (Alickovic et al., 2019). Therefore the EEG data used for decoding underwent no blink rejection or visual evoked potential response pre-processing. The EEG data was bandpassed between [2,32] Hz using EEGLab’s Hamming windowed FIR filter (Delorme & Makeig, 2004; Ciccarelli et al., 2019). The ideally separated talkers’ broadband audio envelopes were extracted using a nonlinear, iterative method (Horwitz-Martin et al., 2016). The bandpassed data and the audio envelopes were then downsampled to 100Hz.

The decoder was implemented using leave-one-trial-out cross validation. The attention decoders were trained using attended and unattended talker envelopes. The switch conditions’ attended talker envelopes had to be constructed out of concatenated talker envelopes from the two talkers. The sustained condition’s attended envelope was composed of a single talker per trial. The ignored talker envelope was constructed using either the switch or sustained condition’s talker envelope method mentioned above. The decoders are talker-invariant; training used a balanced amount of each talker’s data. The decoder used a sliding EEG window of 250 milliseconds (Ciccarelli et al., 2019). Regularization was performed using L2 (ridge regression). Maximizing decoding accuracy was not the focus of the work, we chose to use a fixed regularization of 1e6 for all subjects instead of finding the subject-unique performance-maximizing regularization value via a validation process. The attended talker decision at a given time is determined via Pearson correlation. A Pearson correlation was performed between the candidate speech envelopes, *env*, and the speech envelope prediction, 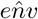. We evaluated the decoder with correlation windows of 10 and 5 seconds. In the results, we present a detailed overview of decoder performance using a length of 5 seconds. The remainder of the analysis uses a 5-second window and using the same window for decoding provides the opportunity to compare features around the switch.

### 2.3. EEG Analysis

#### 2.3.1. Alpha Event Related Spectral Perturbation

EEG alpha was used to quantify measures of listening effort before, during, and after switching between sustained attention. The EEG data was pre-processed differently for this banded analysis than in least-squares. In contrast to single trial decoding, EEG power band analysis is sensitive to blinks. Blink artifacts were removed from the data using independent component analysis (ICA) methods found in the EEGlab toolbox (Delorme & Makeig, 2004). Instead of absolute alpha band power, we computed a measure of relative alpha power in the form of alpha event-related spectral perturbation (ERSP) (Makeig, 1993). Alpha ERSP, *A*(*t, n*), is defined as the difference between the absolute alpha spectral power in a given window and a baseline window, scaled by the baseline window (Eq. 1). Alpha ERSP is a function of both time, *t* and EEG channel, *n*. The spectral density, *P*, was computed across each analysis window using MATLAB’s pwelch method. The absolute alpha power was computed as the sum of squared spectral density values between [8,12] Hz. The baseline window, *B*, indicates that the spectral power was computed across a window between [-25:20] seconds relative to the trial’s timing event (Eq. 1). As a reminder, the timing event is either a switch time or a control for the switch time in the sustained condition. The variable, *t*, indicates that the spectral power was computed on a sliding 5-second window whose latter edge spans [-20:25] seconds relative to the timing event. At 5 seconds for example, ERSP captures the activity between [0,5] seconds proceeding the switch, not just the activity at 5 seconds. ERSP was computed individually for each channel, n, using that channel’s baseline alpha power. We segmented the EEG channels into three sections (frontal, centrotemporal, and parieto-occipital). Then computed the mean ERSP across each of those section’s channels to arrive at a given region’s ERSP measure.

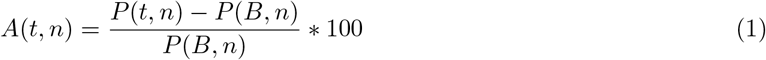

### 2.3.2. Lateralized Alpha Event Related Spectral Perturbation

In addition to listening effort, relative attended and suppressed stimuli locations may also modulate alpha power during auditory attention (Deng et al., 2020). To assess this phenomenon in our data, we extracted an alpha ERSP feature that highlights hemispheric differences in response to attended/ignored talker locations. We used alpha ERSP magnitude from Eq. 1 instead of the individualized measure of peak alpha power magnitude that had been used previously (Deng et al., 2020). For each experimental condition, we computed two time-varying mean alpha ERSP topographies for each subject. One alpha ERSP topography was computed across the 10 trials of a given condition type that were initialized with attention towards the left talker, *A*_L_, and the other with the attention towards the right talker, *A*_R_. The net alpha ERSP, *A*_net_, was defined as the difference in left and right initialized ERSP responses (Eq. 2). *A*_net_ was partitioned into two 20-second segments and used to compute the mean pre-switch and post-switch responses in Eq. 3 and Eq. 4, respectively. The hemispheric difference for a given channel subset, *n*, before the switch, *A*_pre_, was computed using Eq. 5. Similarly, Eq. 6 was used to compute the hemispheric difference for a given channel subset after the switch, *A*_post_. In Eq. 5 and Eq. 6, *N* _L_ and *N* _R_ indicate left and right hemisphere channels for a given channel subset respectively.

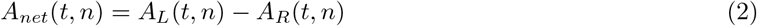

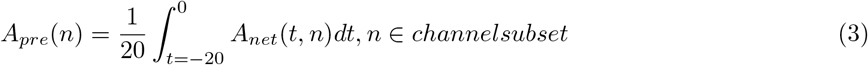

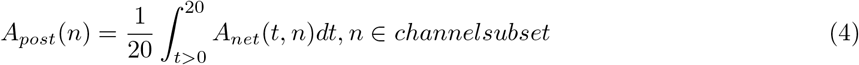

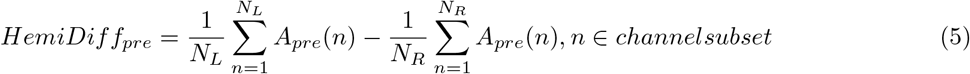

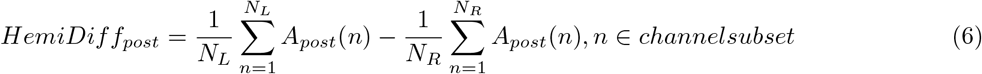

### 2.4. Pupillometry Analysis

We performed peak-based blink detection on the raw pupil diameter data, interpolated data points containing blink artifacts, and smoothed the data using a 1-second median filter. The pupil diameter used for analysis was defined as the average pupil diameter between the left and right pupil channels. We applied the normalization framework from Eq. 1 to pupil diameter (Eq. 7). Pupil dilation was normalized using a trial-by-trial baseline at the onset of the trial since dilation may sensitive to factors unrelated to the experimental task such as engagement, arousal, anxiety, and lighting conditions (van Rij et al., 2019). In Eq. 7, *D*, is the mean pupil diameter in a given 5-second window. Again, *B* and *t* indicate whether mean pupil diameter (MPD) was computed across a baseline window between [-25:20] seconds or a sliding 5-second window whose latter edge spans [-20:25] seconds relative to the time event. Both alpha ERSP and MPD were computed every 10 milliseconds with a 4.99 second analysis window overlap. In addition to the baseline window normalization, we z-scored trial-level ERSP and MPD within each participant, in order to highlight experimental condition differences instead of participant differences.

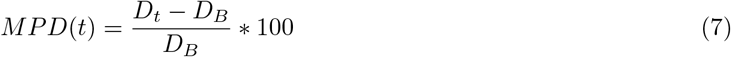

### 2.5. Statistical Analysis

To determine significant differences in decoder, EEG, and pupillometry measures, we ran a two-factor ANOVA tests with experimental condition as a factor and participant ID modeled as a random factor. For each two-factor ANOVA, we performed pairwise t-tests with a Bonferroni correction to illustrate significant differences in population means between experimental conditions. To quantify differences in alpha lateralization in response to attended talker location, we performed a three-factor ANOVA (experimental condition and whether the measure was computed before or after the switch time were treated as factors with participant ID modeled as a random factor). We subsequently performed post-hoc planned comparisons with a Tukey adjustment.

## 3. Results

### 3.1. Attended Talker Comprehension

At the end of each trial, participants answered difficult comprehension questions about the talker(s) they attended to. Listeners answered 120 4-choice comprehension questions with a mean accuracy of 0.56 (SEM = 0.03) which is above chance (0.25). Participants achieved mean comprehension accuracies of 0.58, 0.51, and 0.58 across at-will, directed, and sustained conditions, respectively. A two-factor ANOVA determined that there was a main effect of experimental condition on overall comprehension accuracy [F(2, 18) = 4.247, P = 0.0308]. However, the paired t-tests did not determine that the population means for experimental-condition comprehension accuracy were significantly different. These results suggest that participants performed auditory attention equally as well across the conditions since the three conditions do not have significant differences in their comprehension scores.

### 3.2. Attended Talker Decoding

On a subject-basis, the attention decoders were evaluated using correlation window lengths of 10 and 5 seconds. Decoding accuracy is defined as the fraction of time, the time-varying correlation-based decision vector, *corrDiff*_*acc*_, is correct. *CorrDiff*_*acc*_ is defined as the difference in the time-varying correlation between the decoder output and the two candidate talker envelopes. When performing correlations for accuracy evaluation, the correlation would be computed between the predicted envelope, 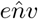, and the true candidate envelopes, *env*_*Att*_ and *env*_*Una*_ (Eq. 8). The grand mean accuracy dropped from 69.3% (SEM = 1.8%)) to 64.1% (SEM = 1.5%) when the correlation window length was halved from 10 to 5 seconds. Moving forward for visualization, the 5-second window was selected since it shared the same duration as the other analyses performed in this study. A two-factor ANOVA found no effect of experimental condition on trial-level decoding accuracy evaluated using the 5-second decision window [F(2,18) = 1.83, P = 0.189] (Figure 2). In addition to *corrDiff*_*acc*_, we computed *corrDiff*_*sw*_, the correlation between 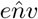 and the initial attended and ignored talker envelopes are represented by *env*_1_ and *env*_2_, respectively (Eq. 9). *CorrDiff*_*sw*_ was computed across experimental conditions. *CorrDiff*_*sw*_ unlike *CorrDiff*_*acc*_ changes sign at the time of the attention switch (Figure 3). *CorrDiff*_*sw*_ crossed zero at 2.31 seconds for the at-will condition and 2.15 seconds for the directed condition. This measure was computed off of the grand mean *CorrDiff*_*sw*_ curve for the two switch conditions. When a listener engaged in a switch in attention, it took approximately half the length of the correlation window for the correlation with the initial talker to weaken below the correlation with the secondary talker (Figure 3).

**Figure 2:**
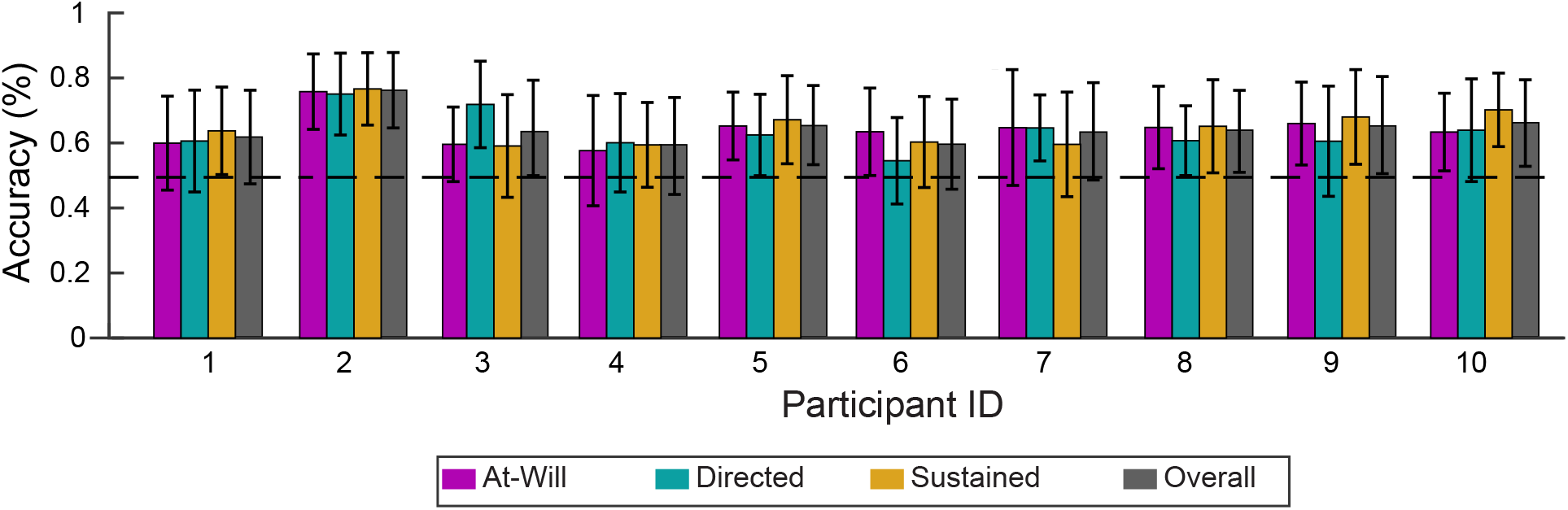
Least-squares attended talker decoding accuracy reported for each participant. Participant-level least-squares attended talker decoding accuracy computed using 5-second correlation window. The grand mean accuracy is 64.1% (SEM = 1.5%). Across experimental conditions; there are no significant differences in accuracy between experimental conditions.

**Figure 3:**
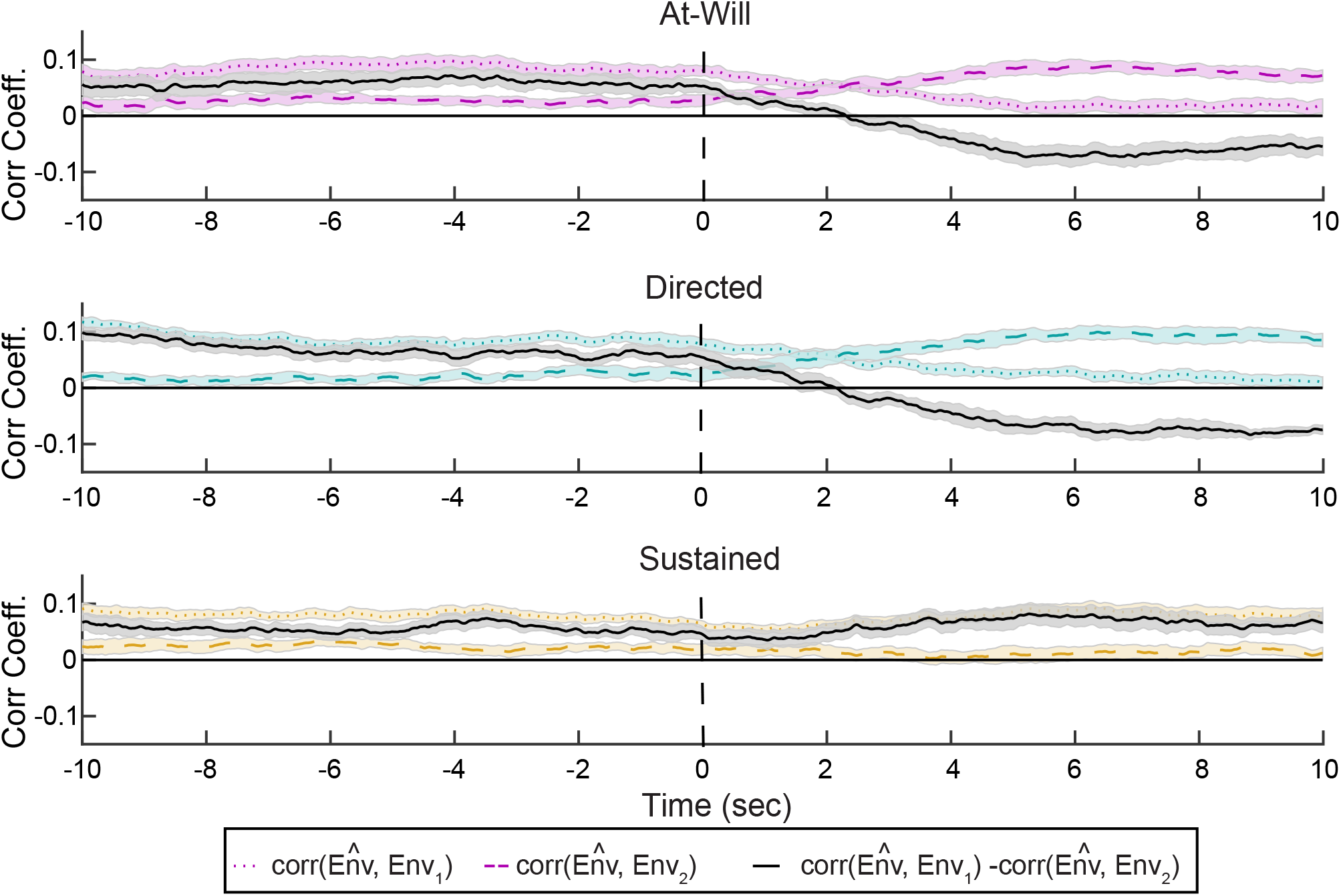
Least-squares decoding illustrates smooth switches in attention. The decoder output was correlated with the trial’s initial attended and unattended talker speech envelopes. In the experimental conditions that contain a switch between talkers (top two panels), the correlation with the attended and unattended talker flip direction. The correlation difference changes sign at a lag of approximately half the correlation window.

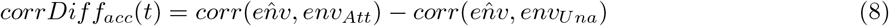

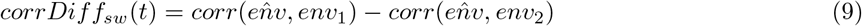

### 3.3. Event Related Spectral Perturbation and Mean Pupil Diameter

Around the normalized time of zero seconds, grand-mean alpha ERSP topographies demonstrate differences in the trials that contain a switch in contrast to the sustained condition (Figure 4). At 5 seconds, the at-will and directed alpha ERSP responses were globally weaker in magnitude than the sustained condition. There were weak main effects of experimental condition on frontal alpha ERSP [F(2, 18) = 3.201, P = 0.0646] and parieto-occipital alpha ERSP [F(2, 18) = 3.426, P = 0.0549]. There was a strong main effect of experimental condition on centrotemporal alpha ERSP [F(2, 18) = 7.473, P = 0.00434]. Time-varying grand-mean centrotemporal alpha ERSP and MPD (with standard error of the mean error bars) illustrate these trial differences at a finer temporal resolution (Figure 5). Both alpha ERSP and MPD are indistinguishable across experimental conditions before time zero, likely because the attention task is similar across the trials in that time region. At zero seconds, alpha ERSP and MPD magnitudes are stacked in value in the order of task difficulty. Paired t-tests found that the centrotemporal alpha ERSP population means are different between the sustained condition and the two switch conditions at a significance level of 0.042 (at-will) and 0.018 (directed) (Figure 6A). There was a strong main effect of experimental condition on the 5-second MPD value [F(2, 18) = 9.159, P = 0.0018] as well. Paired t-tests also found that MPD population means are different between the sustained condition and the two switch conditions at a significance level of .017 (at-will) and 0.021 (directed) (Figure 6B).

**Figure 4:**
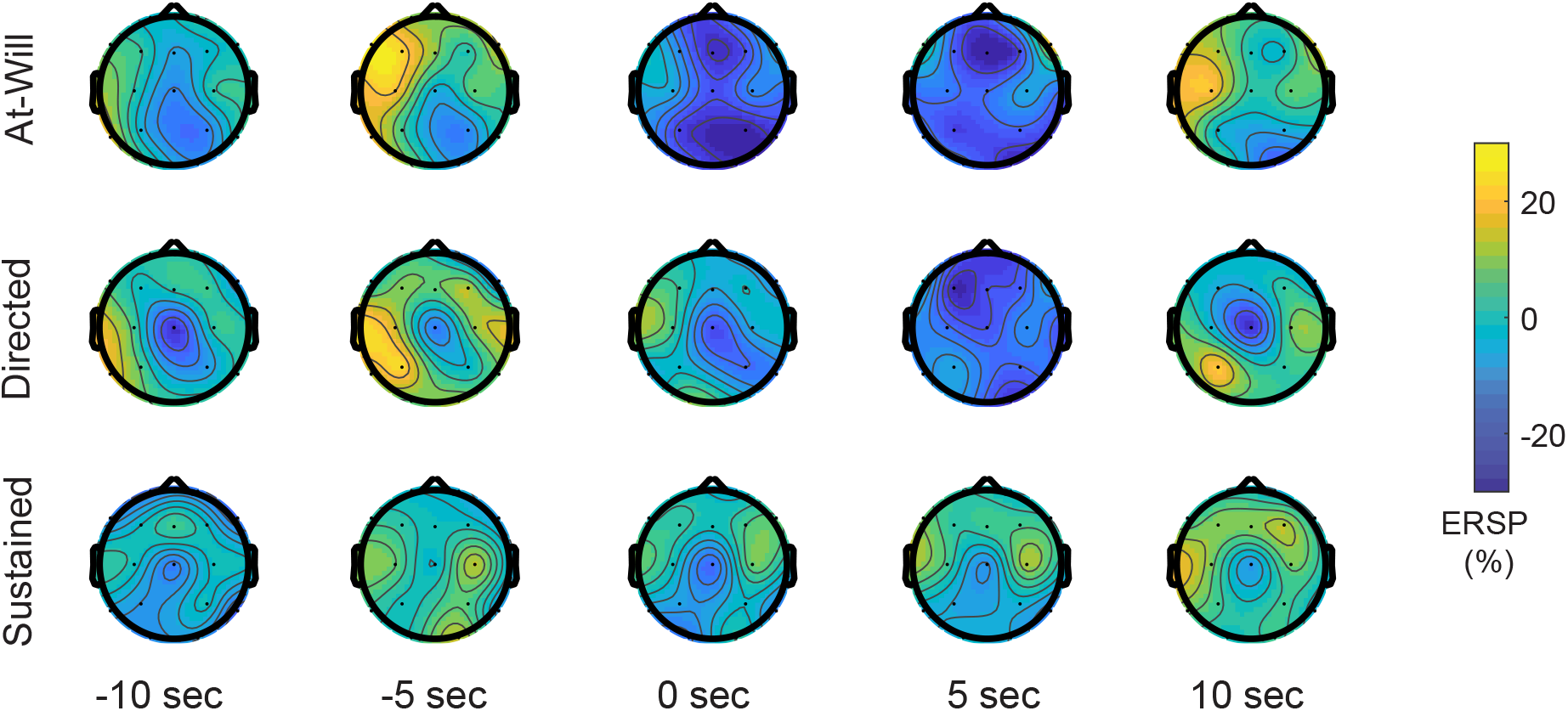
Grand mean alpha event related spectral perturbation topographies. Grand-mean z-scored alpha ERSP topography sampled at time points before, during, and after the switch time for each experimental condition. Alpha ERSP was computed using a sliding 5-second window of data, therefore the sampled topographies shown capture activity from the preceding 5 seconds of time. At 5 seconds, the switch experimental condition topographies have weaker alpha ERSP magnitudes than the sustained experimental condition.

**Figure 5:**
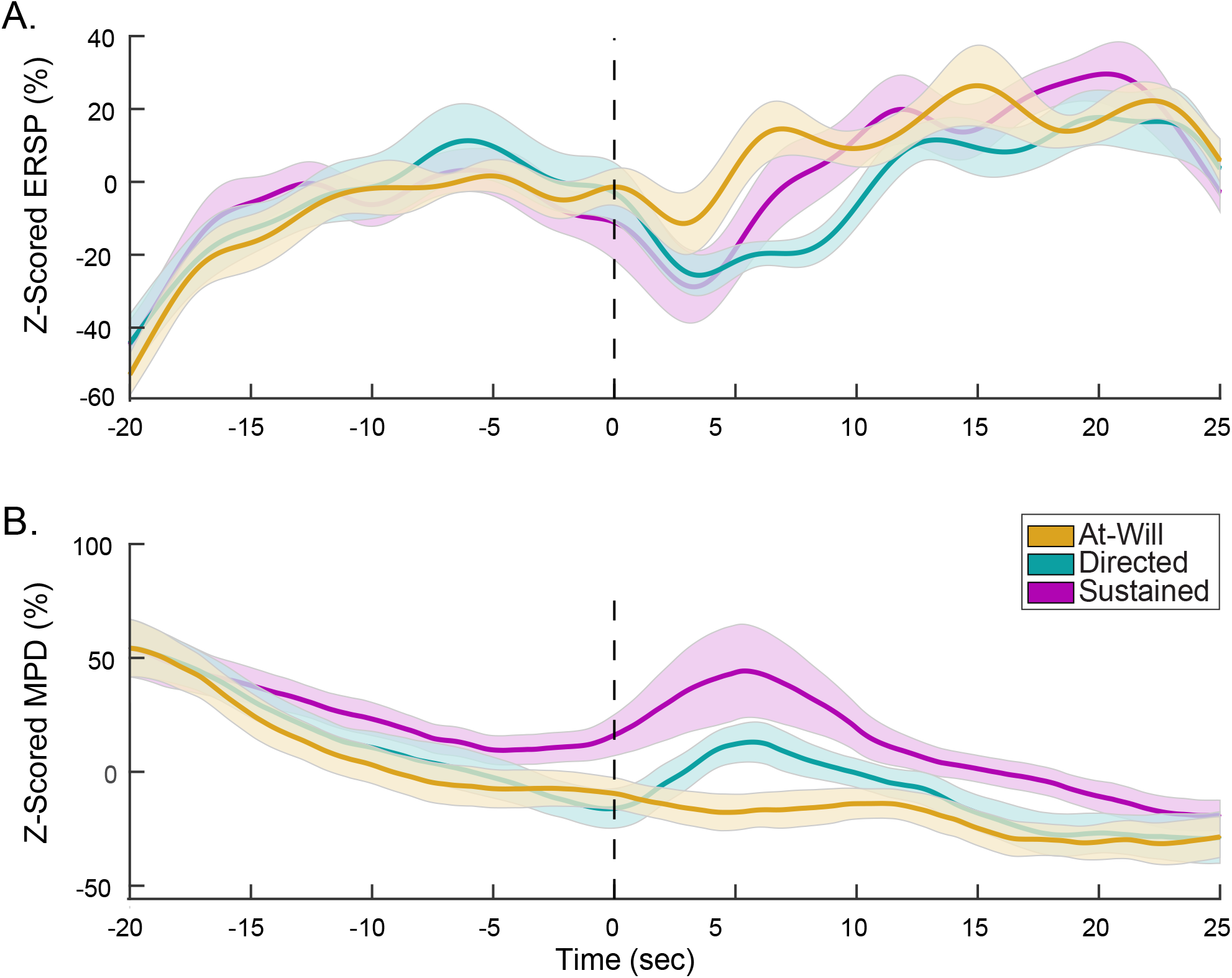
Centrotemporal alpha event related spectral perturbation and mean pupil diameter over time. (A) Grand mean (and standard error of the mean) centrotemporal alpha event related spectral perturbation (ERSP) and mean pupil diameter (MPD) response over the course of each experimental condition. Centrotemporal alpha ERSP and MPD trend in opposite directions over the course of the trial. Around the switch time, ERSP and MPD capture experimental condition differences.

**Figure 6:**
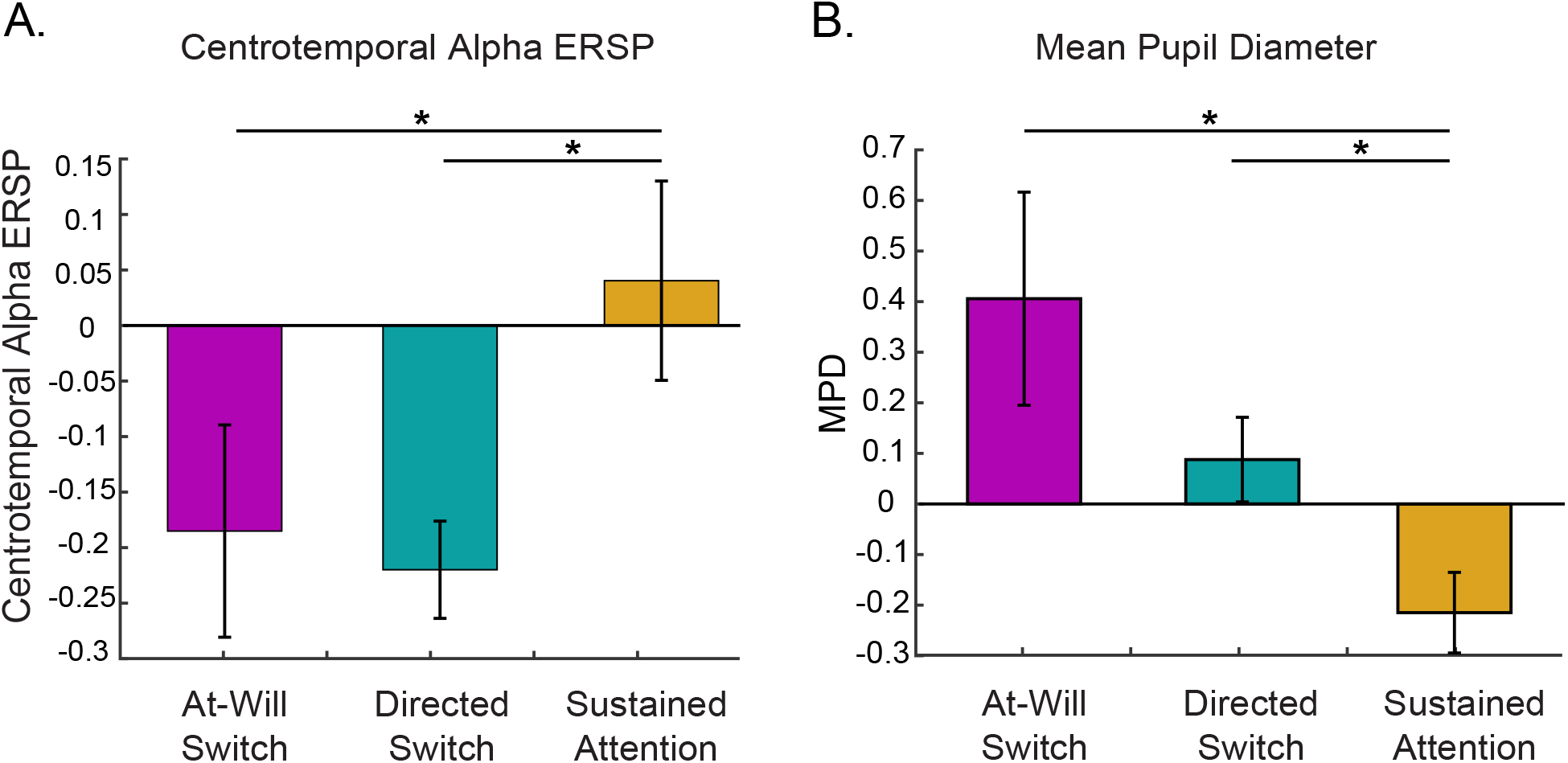
Centrotemporal alpha event related spectral perturbation and mean pupil diameter at 5 seconds after switch. In the 5 seconds following the switch, both centrotemporal alpha ERSP and mean pupil diameter responses to the sustained experimental condition are significantly different from the switch experimental conditions.

### 3.4. Alpha Lateralization

The topographic distribution of alpha power relative to the attended and unattended talker locations illustrate differences between experimental conditions (Figure 7). The grand-mean net alpha ERSP topographies before and after the switch (A_pre_ and A_post_) show that switch trials reflect a change in the dominant alpha hemisphere in the centrotemporal region (Figure 7A). For all three experimental conditions, A_pre_’s left hemisphere is ipsilateral to attended talker in Eq. 2’s leading term. A_post_’s ipsilateral/contralateral hemisphere demarcation is dependent on the experimental condition. For the sustained condition, the initial attended talker remains the leading term in Eq. 2. For the two switch experimental conditions, the initial attended and ignored talkers switch roles in the latter half of the trial after a switch. If a switch occurred, Eq. 2’s leading term contains the response to right talker attention instead of left talker attention. This reshuffling of Eq. 2’s leading terms changes which hemisphere is considered A_post_’s ipsilateral hemisphere.

**Figure 7:**
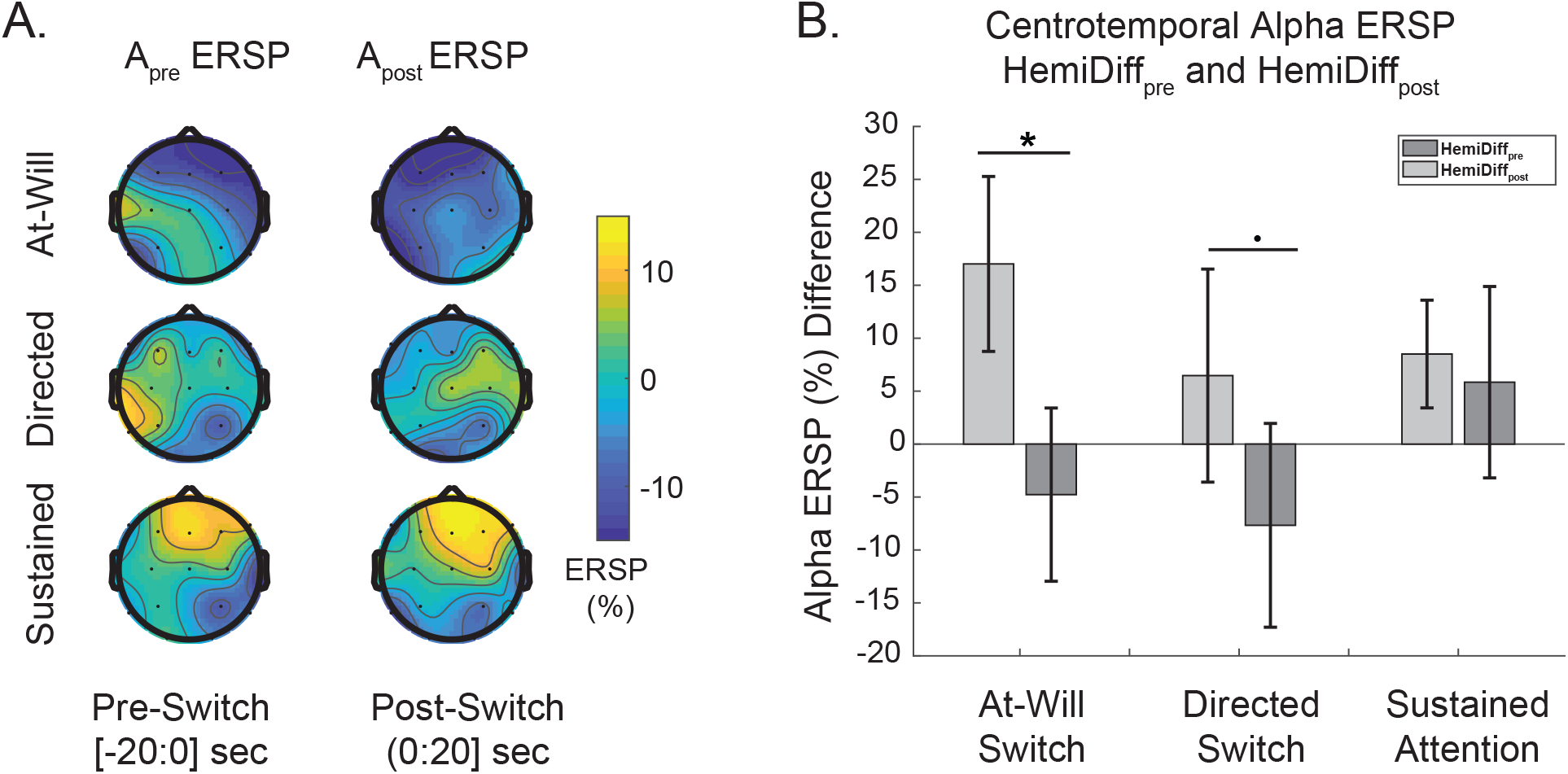
Alpha event related spectral perturbation lateralization. (A) The net alpha ERSP topography averaged across 20 seconds before and after the switch. In the switch experimental conditions, the hemisphere with the greater centrotemporal alpha magnitude switches hemispheres in keeping with the hemisphere that is ipsilateral to the attended talker for the given segment of time. In the sustained experimental condition, centrotemporal alpha magnitude remains on the same side since the attended talker location did not change at time 0 seconds. (B) Together, these responses support alpha ERSP lateralization ipsilateral to the attended talker location.

These hemispheric differences are further quantified by HemiDiff_pre_ and HemiDiff_post_ (Eq. 5 and Eq. 6). This measure shows an alpha dominant hemisphere shift occurs for the switch trials and not the sustained trials (Figure 7B). We performed a three-way repeated-measures ANOVA (experimental condition by whether the measure was computed before or after the switch time) with subject ID modeled as a random factor. There was no main effect of experimental condition on centrotemporal alpha HemiDiff [F(2, 18) = 0.511, P = 0.609]. There was a strong effect of pre/post switch time on centrotemporal alpha HemiDiff [F(1, 9) = 23.01 P = 0.000978]. There was a weak interaction between the experimental condition and pre/post switch time on centrotemporal alpha HemiDiff [F(2,18) = 3.227, P = 0.0634]. The planned comparison found that within switch conditions, the centrotemporal alpha HemiDiff_pre_ and HemiDiff_post_ population means are different at uncorrected p-values of 0.0335 (at-will) and 0.0885 (directed), respectively. Additionally, for the sustained experimental condition, the centrotemporal alpha HemiDiff_pre_ and HemiDiff_post_ population means are not significantly different. The planned comparison results support attended/unattended talker location modulated alpha lateralization.

## 4. Discussion

We studied endogenous attention switching in the context of developing decoding algorithms that can be used in natural, every-day multi-talker listening environments. Our experimental protocol allowed listeners to endogenously switch attention between continuous speech sources while their effort was characterized through EEG and pupillometry measurements. In addition to effort, we detected two types of endogenous attention switches using both talkers’ speech envelopes and spatial locations. While this decoding result is not the first to demonstrate endogenous attention switch decoding (Miran et al., 2018, 2020), to the best of our knowledge, it is the first study to decode multi-talker continuous speech without the potential confound of sensorimotor planning. We also characterize the effort involved with attention switching between speech sources. This builds upon effort measures associated with sustained attention between competing speech sources (Seifi Ala et al., 2020) and attention switching between a pairs of competing alphabetic characters (McCloy et al., 2017). EEG alpha power and pupil diameter measures indicated that the effort associated with performing attention switches was greater than our sustained-attention condition. Listener centrotemporal alpha power was also found to be modulated by the relative spatial locations of the stimuli. Our decoding results highlight latencies inherent in speech-feature decoding. Our EEG and pupil diameter findings support leveraging attention-switch decoding and other non-speech features for improving the accuracy and decreasing the decision latency involved with cognitively-controlled hearing aids.

### 4.1. Switching Latency of Envelope-Based Attention Decoding

Listeners who struggle with speech understanding in multi-talker scenes would significantly benefit from enhancement that instantaneously cues on the talker they wish to attend. For practical applications, decoding algorithms must operate in a causal manner, incrementally producing a decoding decision from a given window of previous data. When there is a switch in attention, this analysis window length translates into a decoding latency. Although the 5-second correlation window would produce a faster detection of a switch than a 10 second correlation window, it is at a cost. The 5-second correlation decision produced noisier predictions over time and reduced decoding accuracy by 5.2% when compared to the 10-second window. This trade-off was systematically studied with a linear model on another data set that contained simulated attention switches and optimal accuracies of 62% and 68% were achieved using an evaluation window of 2.54 and 11.28 seconds (Geirnaert et al., 2019).

Our study in contrast, evaluates performance on EEG data that contains real human switches in attention and confirms that switches can be detected in approximately half the decision window size using a standard least-squares decoding method. We originally hypothesized that when an attention switch occurs, there is a measurable latency associated with the time it takes for the listener to go from attending to one source to another. While the decoding lag defines the fastest the decoder can detect an attended talker change, it assumes a negligible human switching delay. In both 10 and 5 second evaluations of our decoder, we observed the decision vector (Eq. 9) change sign at a lag of half the respective correlation window length, indicating a switch in the listener’s attended talker. For the 5-second correlation window length, mean switch time was 2.31 and 2.15 respectively for our at-will and directed experimental conditions. Since the decoded switch time was less than half the window size, this indicates that listeners were potentially switching slightly before the reported switch time, and that there was no additional lag associated with attention switching. A state-modeling algorithm found algorithmic delays of 1.9, 1.75, and 1.5 seconds for simulated switch data, real switches in EEG, and real switches in MEG, respectively (Miran et al., 2018). The decoding lag in our data set and others, demonstrates the need for further inquiry into alternative decoding features such as expended effort, laterality due to spatial cues, pupillometry, and eye-gaze. Supplementing envelope-decoding with other features may further reduce the algorithmic switching time for attention decoding.

### 4.2. Increased listening effort is associated with auditory attention switching

Around the switch time, both alpha ERSP and MPD demonstrated differences in value for the switch and sustained conditions. These magnitude differences were superimposed on slow alpha ERSP and MPD trends in opposite directions over the course of the trial. This slow increase in alpha ERSP and decrease in MPD was also present in previous work (Seifi Ala et al., 2020). Its not clear whether these changes in ERSP and MPD are related to a change in effort and may just be a physiological adaptation. Our results suggest that the effort required to switch attention was greater than the effort required to sustain attention. The raw alpha ERSP and MPD magnitudes observed in Figure 5 reflect both the effort due to switching and higher-order cognitive tasks (in-the-moment decision making and time memorization), depending on the experimental condition instructions. In our experiment, both the at-will and sustained conditions involve decision making and time memorization and only differ in whether an attention switch occurs. Therefore the differences in switch and sustained condition ERSP and MPD measures are due to the effort required to implement the switch in attention.

Our alpha power results are consistent with previous sustained attention effort characterization (Seifi Ala et al., 2020). Where a lower magnitude alpha ERSP was associated with greater listening effort, we found a similar result in our most difficult experimental condition (at-will). As expended effort increases, cortical networks activate, resulting in decreased cortical synchrony and decreased alpha ERSP (Seifi Ala et al., 2020; Pfurtscheller, 2001; Jensen & Mazaheri, 2010). On the other hand, our alpha ERSP findings differ from previous findings in the specific EEG channel subset where the alpha ERSP effect was found. Significance was found in the parietal channels (Seifi Ala et al., 2020) while we found significance in the centrotemporal channels. This discrepancy may be due to a difference in participant age and hearing-aid usage (Seifi Ala et al., 2020). We believe EEG alpha power should be used in addition to other factors such as attended stimulus entrainment and pupil diameter measures due to the band’s ability to be modulated by other factors. Our results show that pupil diameter increases during our complex attention switching tasks in manner that is consistent with previous pupil diameter measures performed during an exogenous attention switch between competing alphabetic character pairs (McCloy et al., 2017).

In addition to understanding the effort associated with attention switching, pupil diameter measured throughout the entire 60-minute collection can be leveraged to determine the impact a listener’s effort has on decoding accuracy. In future studies, pupil diameter can be used as a measure of fatigue over the course of long stretches of effortful listening and to determine auditory training’s efficacy in reducing such fatigue (Pichora-Fuller et al., 2016). These attention switching conditions could be implemented in clinic to gauge listener effort when performing auditory attention between stimuli with low speech intelligibility (Winn et al., 2018; Zekveld et al., 2018; Pichora-Fuller et al., 2016; Paul et al., 2021). These measures could help gain insight on an individual’s fatigue associated with everyday difficult listening conditions out side the clinic as well.

### 4.3. EEG alpha power is lateralized by attentional spatial cues

Our results confirm the hypothesis of the at-will and directed switch conditions having significant differences in the hemispheric difference measure before and after the switch. Several prior studies have suggested that the spatial location of acoustic stimuli lateralizes alpha power during an attention task (Bonnefond & Jensen, 2012; Bednar & Lalor, 2018; Weisz et al., 2011; Deng et al., 2020). Stimulus suppression has been shown to increase alpha in the hemisphere ipsilateral to the attended talker in an attention task of competing syllables (Deng et al., 2020). We found greater alpha magnitude in the hemisphere ipsilateral to the attended talker as well (Figure 7). Recall that this alpha power difference between hemispheres was quantified as a measure of hemispheric difference as shown in Eq. 5 and Eq. 6. A strong effect of pre/post switch time was found to impact the hemispheric difference measure. This factor is likely capturing that before the switch, all experimental conditions have a lower mean alpha ERSP than after the switch Figure 5. However, in addition to that effect, we also found a weak, yet still present, interaction between the experimental condition and whether the measure was computed before or after the switch on the hemispheric difference measure [P*<*0.1]. This interaction captures the fact that the hemispheric difference measure embeds trial dependent talker location movements (Eq. 2-Eq. 6). The uncorrected planned comparisons also support relative stimuli location alpha modulation. The hemispheric difference measure before and after a switch is expected for the switch conditions because the hemisphere ipsilateral to the leading term in *HemiDiff*_*pre*_ and *HemiDiff*_*post*_, switches when the listeners switch attention (Eq. 5,Eq. 6). In contrast, our sustained experimental condition did not have the listener switch attended talker locations and our results show that there was no statistical difference between the sustained condition’s alpha hemispheric difference before and after the switch.

Alpha lateralization was significant in the centrotemporal region. This alpha lateralization result is consistent with a previous finding, although the cortical region differs slightly from the parieto-occipital region previously identified (Deng et al., 2020). One explanation for this difference is that our task was significantly longer and more complex. Another possibility could be related to the fact that we computed alpha ERSP instead of individualized peak alpha magnitude for our alpha lateralization measure. Regardless, our results further support that even with a demanding task of attention between continuous speech stimuli, alpha lateralization effects are present. Although there is evidence of spatially modulated alpha power, this cue is limited for single-trial decoding use due to the window length over which the feature was computed for significance. Although this effect exists, better features may exist for leveraging spatial cues for decoding. For example, a decoding method that used common spatial pattern filters to determine directional focus without the use of speech-features, performed at an accuracy of 80% and window length of 1 second (Geirnaert et al., 2020).

### 4.4. Leveraging attention switches for a cognitively-controlled hearing aid

Cognitively-controlled hearing aids have the capacity to improve the listener experience in cluttered environments through listener-steered speech enhancement (Geirnaert et al., 2021). Understanding endogenous switching may speed attention decoding through intended attention identification throughout switching before a new talker is fully attended to. Speech-feature based decoding relies on the attended speech being encoded in the listener’s cortical signals. It remains unknown how these speech-feature based algorithms would work on real attention switches in individuals with hearing impairment (Decruy et al., 2020; Van Canneyt et al., 2021). On the other hand, sensing effort expended in an attempt to attend to a new source and ignore another, could be leveraged to help decode intent in this situation. It is probable that the attention processes involved with an endogenous switch may begin to show themselves in cortical signals earlier than an exogenous switch due to the decision-making and planning involved. Therefore, supplementing speech-feature based decoding with features that are directly related to switches in auditory attention, may result in decreased decoding lag and increased accuracy. The neural and pupil diameter markers associated with switching listening effort, as shown in our results, could potentially be leveraged as one of these features.

We believe this work further supports exploring non-acoustic, multi-modal features for attention decoding. Our results demonstrated that speech-feature based decoding still functions in the presence of additional higher-order cortical tasks, indicating that non-speech features have promise to be fused with speech-features for robust multi-cue feature decoding. This work did not focus on maximizing decoding accuracy nor minimizing the switch detection lag but future work could aim to use these additional features as part of decoding models. Specifically, alpha ERSP and pupil diameter features may be relevant since they both began to change their slope behavior slightly before or at the time listeners reported their switch. Individuals naturally also use both auditory and visual attention in a multi-talker listening task, therefore eye gaze can also be pursued as a non-covert feature for auditory attention decoding (O’Sullivan et al., 2019; Best et al., 2017; Favre-Felix et al., 2018).

## 5. Conclusion

In this study, we characterized the effort associated with endogenous auditory attention switching using both cortical and pupil diameter measures. Decoding real endogenous switches in attention illustrated the problematic lag associated with decoding methods that rely on attended talker speech features. Alpha ERSP and MPD measures of effort were sensitive to endogenous switching of auditory attention. Our effort-based features have a potential application in a multi-modal, multi-feature decoding algorithm for use in a cognitively-controlled hearing aid. Both effort features hold promise in being quick to reflect the onset of switching while being stable in their time course, potentially leading to a shorter lag in switch detection. The study’s effortful attention switching tasks may also apply to the development of objective neural markers of listening effort that are intended for clinical use (Paul et al., 2021; Pichora-Fuller et al., 2016; Zekveld et al., 2018). One last application of these switching effort measures is in the field of attention disorders and development (Hanania & Smith, 2010). Characterizing auditory attention across populations and within individuals is important to pursue in combination with developing effort-based features for decoding. In addition to clinical hearing ability (Vanthornhout et al., 2018; Fuglsang et al., 2020; Decruy et al., 2020), expended cognitive effort during listening may significantly impact an individuals auditory attention decoding accuracy. Cognitive-controlled hearing-aid technology can leverage listener effort in many ways. Decoding algorithm speed and accuracy, listener benefit due to enhancement, and efficacy of auditory training can all utilize measures of effort.

## Author Contributions

SH: Conceptualization, Methodology, Formal Analysis, Software, Visualization, Writing-Original Draft Preparation, Writing-Review and Editing. HMR: Pupillometry Analysis, Writing-Review and Editing. TFQ: Conceptualization, Writing-Review and Editing. CJS: Conceptualization, Funding Acquisition, Methodology, Project Administration, Writing-Review and Editing.

## 6. Acknowledgments

The authors would like to acknowledge the organizers the 2019 NSF-funded Neuromorphic Cognition Engineering Workshop in Telluride, CO, USA where this protocol was conceived and pilot data was collected. We thank Edmund Lalor for providing the stimulus and comprehension questions, and Pablo Ripolles for helping collect the pilot data. We would also like to thank workshop participants who participated in discussions regarding this protocol, including Malcolm Slaney, Gregory Ciccarelli, Edmund Lalor, Alain de Cheveigne, Behtash Babadi, Nima Mesgarani, and Tom Francart.

## 7. Supplemental Materials

We computed ERSP measures across the three lowest cortical power bands (alpha, theta, and delta) and across the frontal, centrotemporal, and parieto-occipital EEG channel subsets. In addition to the centrotemporal alpha ERSP and lateralization findings described in the main text, we also wish to report the following additional statistically significant ERSP results.

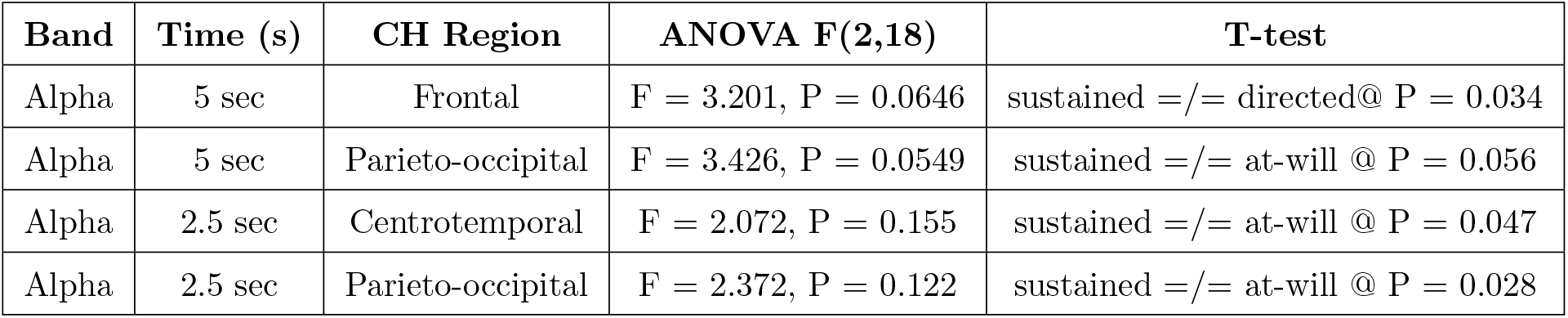
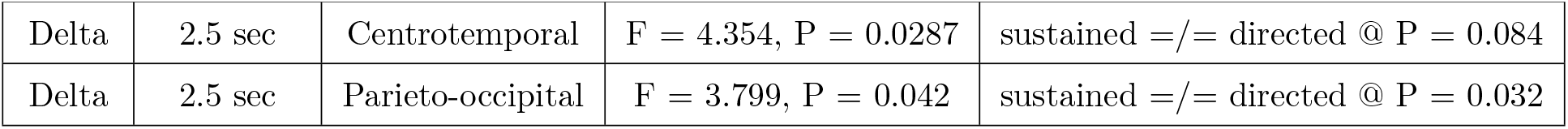

